# Major Update and Improved Validation Functionality in the mwtab Python Library and the Metabolomics Workbench File Status Website

**DOI:** 10.64898/2025.12.19.695605

**Authors:** P. Travis Thompson, Hunter N.B. Moseley

## Abstract

The Metabolomics Workbench (MW) is a public scientific data repository consisting of experimental data and metadata from metabolomics studies collected with mass spectroscopy (MS) and nuclear magnetic resonance (NMR) analyses. Although not as rapidly as in the past, MW has steadily evolved; updating its mwTab and JSON deposition text file formats and its web-based infrastructure. However, the growth of MW has been exponential since its inception in 2013 and continues to be exponential, with the number of datasets hosted on the repository increasing by 50% since April 2024. As part of regular maintenance to keep up with changes to the mwTab file format and an earnest effort to use MW datasets in meta-analyses, the mwtab Python package has been updated. Updates include better error handling for batch processing, better parsing to read more files without error, and extensive improvements to the validation capabilities of the package. These updates also required our mwFileStatusWebsite to be updated and improved. We used the enhanced validation features of the mwtab package to evaluate all available datasets in MW to facilitate improved curation, FAIRness of the repository, and reuse for meta-analyses. Version 2.0.0 of the mwtab Python package is now officially released and freely available on GitHub and the Python Package Index (PyPI) under a Clear Berkeley Software Distribution (BSD) license with documentation available on GitHub. The updated mwFileStatusWebsite is also officially in its 2.0.0 version and is still available at https://moseleybioinformaticslab.github.io/mwFileStatusWebsite/.

## 1. Introduction

The National Institutes of Health (NIH) Reform Act of 2006 established the NIH Common Fund to support cross-cutting, trans-NIH programs. The Common Fund’s Metabolomics Program, started in 2012 and supported until 2022, established the Metabolomics Common Fund’s National Metabolomics Data Repository known as the Metabolomics Workbench (MW) as a longstanding, public repository for national and international metabolomics data and metadata [1]. This repository consists of experimental data and related metadata for metabolomics studies collected with mass spectroscopy (MS) and nuclear magnetic resonance (NMR) analytical platforms. MW is organized into specific projects, studies, and analyses (experiments). Each analysis can be accessed and downloaded in mwTab tabular [1] or JavaScript Object Notation (JSON) formatted files [2,3], both of which are text-based. MW offers a web interface to search, access, deposit, or download organized metabolomics data and metadata. Additionally, MW offers a REpresentational State Transfer (REST) interface to download and view (meta)data [4]. Since MW’s establishment in 2013, the repository has grown exponentially, roughly doubling every two to three years. As of October 22, 2025, MW contained a total of 2466 projects, 3795 studies, and 6125 analyses.

The mwtab Python package was originally released in 2017 to facilitate Python programmatic access to mwTab formatted files from MW [5]. This package also had basic facilities for creating mwTab formatted files to facilitate more automated deposition processes. We then released the first major 1.0.1 version written in Python 3, which dramatically increased the functionality of the mwtab package [6]. Release 1.0.1 included updates to reflect the evolving changes in the mwTab format specification, incorporated methods that utilized the MW’s REST interface, and provided a range of new validation functionality for quality control and curation purposes. However, the overall goal of the mwtab Python package is to promote the FAIR data principles of findability, accessibility, interoperability, and reusability [7,8] with respect to MW. Now we are releasing the next major 2.0.0 version, which contains updates for every module in the package and adds 2 new modules. A summary of the changes to each module are shown in Table 1.

**Table 1.** Summary of changes made to each module in the new 2.0.0 release.

| Modules | Changes |
| --- | --- |
| cli | Refactored for DRY (Don't Repeat Yourself) purposes.<br>Added error recovery for many commands. |
| converter | Added error recovery when converting multiple files.<br>Fixed write out errors due to non-ASCII characters. |
| fileio | Refactored for DRY purposes.<br>Added error recovery when reading and writing multiple files.<br>Fixed write out errors due to non-ASCII characters.<br>File paths that don't exist will be created instead of raising an error. |
| mwextract | Added error recovery when operating on multiple files.<br>File paths that don't exist will be created instead of raising an error. |
| mwrest | Added error recovery when operating on multiple files. |
| mwschema | Completely refactored to use the jsonschema package instead of the schema package.<br>Added more validations, particularly on values of some attributes. |
| tokenizer | Improved parsing, making some formerly unreadable files readable. |
| mwtab | Fixed round trip errors by adding support for duplicate keys.<br>Key/block order is now always the same when writing out.<br>Added convenience properties for attributes in the METABOLOMICS WORKBENCH header.<br>Added convenience methods to convert the tabular sections of the mwTab format to pandas DataFrames and vice versa.<br>Improved file parsing by taking into account some common errors in the mwTab files. |
| validator | Added many new validations.<br>Improved issue printing by standardizing formatting and greatly reducing printing of spurious issues.<br>Added a new JSON output with all validation messages tagged with 'format', 'value', or 'consistency'. |
| duplicates_dict | A new module that contains the DuplicatesDict class used to support reading and writing duplicate keys. |
| metadata_column_matching | A new module for matching metadata columns by names and values.<br>Contains regular expressions, functions, and classes used for matching metadata columns.<br>Contains the "column_finders" dictionary of prebuilt ColumnFinder objects used to match the most common metadata columns in the MW repository. |

The most significant changes in the new mwtab release are in the validations. Specifically, many more validations have been added, and they have been classified based on the nature of the issue with the mwTab file. These changes necessitated an update to another related package and website that we maintain, the mwFileStatusWebsite [9]. This package is also being released to a new 2.0.0 version. The most significant changes to the website are adding a new status, “Warnings Only”, recoloring some of the statuses, and adding a new “Validation Error Type” table that breaks down the validation errors into the new classifications and links to new pages that filter the files based on those classifications. These changes are best illustrated in Figure 1.

**Figure 1.**
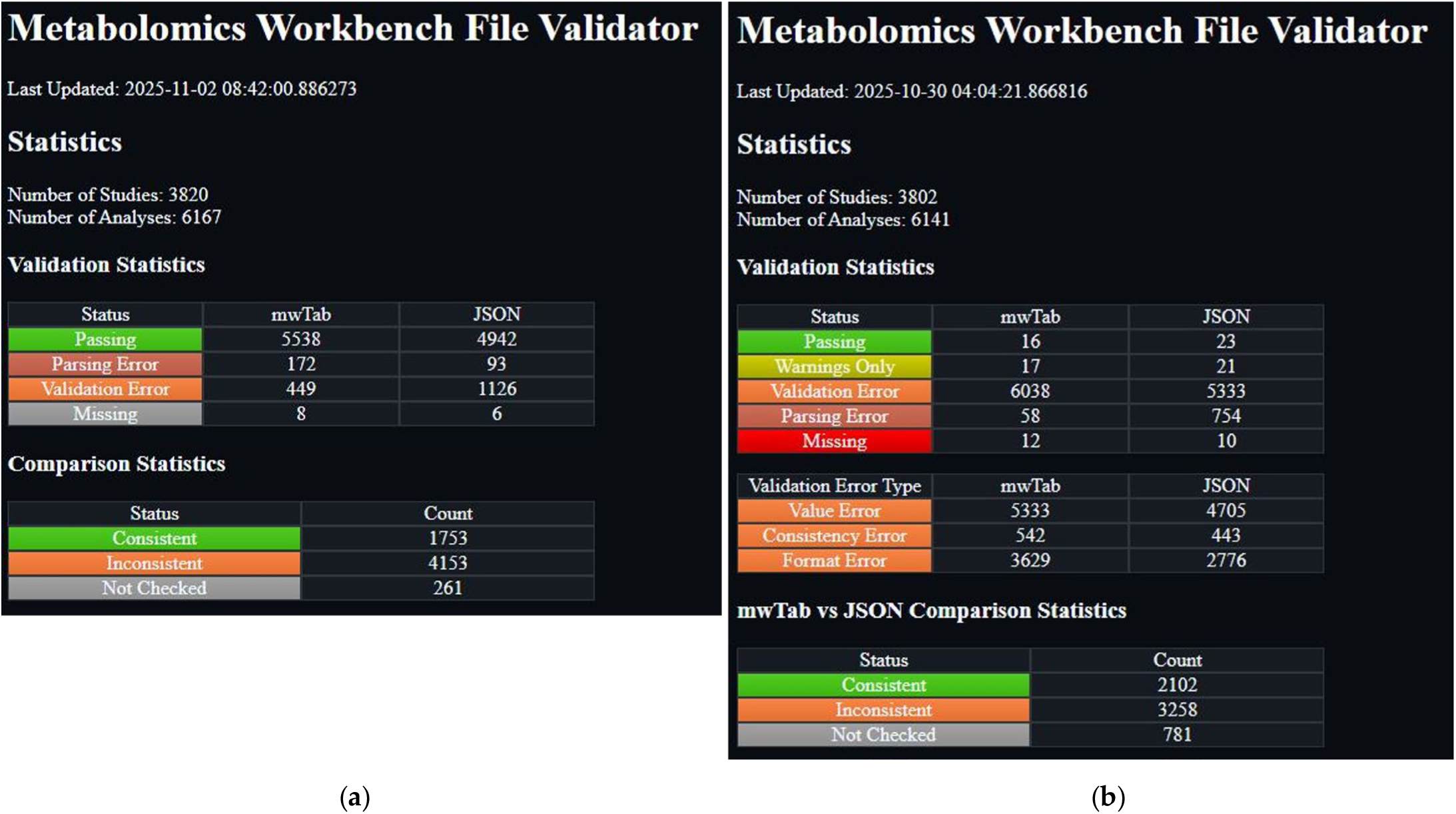
Screenshots from the previous and new versions of the mwFileStatusWebsite: (**a**) Screenshot of the previous version of the mwFileStatusWebsite; (**b**) Screenshot of the current version of the mwFileStatusWebsite.

## 2. Materials and Methods

### 2.1. Updates to the mwTab Format

The mwTab format specification was last updated on 01 August 2023. The updated mwTab format specification is available on the MW website [10].

The mwTab format has remained mostly the same except for a few additions. These additions to the mwTab format include: (1) changing 5 attributes in the CHROMATOGRAHPY section from optional to required, (2) deprecating the NMR_BINNED_DATA section in favor of putting that data in a separate file indicated by a NMR(MS)_RESULTS_FILE attribute in the NM(MS) section, and (3) similarly allowing and preferring unassigned mass spec measurements to go in a separate file indicated by the MS_RESULTS_FILE attribute in the MS section. Many of these items are not explicitly stated as being updates to the mwTab format, but as of November 2025 are present in a significant number of analyses. Tracking updates to the mwTab specification is not easy for 3 reasons. First, older versions of the specification are not readily available from MW website, only the most recent one. Second, although there is a VERSION attribute in the mwTab file header, it is not effectively used. All analyses just have the value of ‘1’ for this attribute, meaning that the version of the specification used to generate the file is unknown. Third, not all changes are explicitly stated in the specification file and instead must be inferred through observation of the files or use of the online submission system. When using the web forms to try and submit to MW, there are differences in what the forms require vs what is stated in the mwTab specification.

### 2.2. mwtab Package Implementation

The mwtab Python package previously consisted of nine modules: mwtab.py, tokenizer.py, fileio.py, converter.py, mwschema.py, validator.py, mwextract.py, mwrest.py, and cli.py. In the 2.0.0 release, changes were made to all of those modules, and two additional modules, metadata_column_matching.py and duplicates_dict.py, were added to the package. The updates to the package modules largely fall into three main categories: (1) updates to improve error handling when operating on multiple files, (2) updates to improve the main functionality of the module or fix bugs, and (3) a great expansion of the mwTab format and metadata validation capabilities along with the addition of some data validation capabilities.

The cli.py, converter.py, fileio.py, mwextract.py, and mwrest.py modules were all upgraded with code to recover from an unexpected error when processing one file out of multiple files. Previously, for some functions, if an unexpected error was encountered while processing one file in a batch of files, the overall batch process would error and the remaining files would not be processed. Many of the aforementioned unexpected errors were also investigated and the code was improved to handle those situations. For example, if an output path was specified, but one or more of the directories did not exist, this would cause an error. Code was added to check for this situation and create the directories that do not exist. The tokenizer.py module also underwent this improvement process but did not need the upgrade to recover when batch processing, because it does not do batch processing.

The mwtab.py module received significant upgrades and additional functionality. The most significant improvement is the ability to handle duplicate keys. Round trip testing refers to using the package to read in a file and then write it back out with no information loss. When performing this testing, it was discovered that there was information loss for some files and it was due to some parts of the file having duplicate names. For example, if two columns in the METABOLITE_DATA or METABOLITE sections have the same name, then only the last column with that name would be read in. This is due to Python dictionaries not being allowed to have a key that points to two different values, so when the second column was added to the same dictionary that the first was already in, it overwrote the first column’s data. This was a difficult error to find and fix. Difficult to find because there is no warning about this duplicate key from Python’s built-in json package used to read in the JSON version of mwTab files. Difficult to fix because although duplicate keys are technically allowed in JSON, they are HIGHLY discouraged and therefore such a rare occurrence that there is no standard way in Python to read and write them out. The new module, duplicates_dict.py, was created to handle this situation. It contains a single class, DuplicatesDict, that is used instead of the regular Python dictionary to be able to read in and write out duplicate keys in JSON. Other improvements to the mwtab.py module include working hand-in-hand with the tokenizer.py module to improve parsing, adding convenience properties and methods when working with the API, and improving writing out files by setting the section/key order to a predefined order so all files write out with the same section/key order.

The validator.py and mwschema.py modules work together and received the most significant changes. The mwschema.py module was completely rewritten using the jsonschema Python package [11,12] instead of the schema Python package. This was to enable both better error messaging and better validation. The jsonschema package has better support in both of these areas. The schemas were also improved to validate the values of some attributes with a more constrained value space, such as STUDY_ID. Several functions were also created to help construct the regular expressions used for value validation and to help construct the custom error messages for those attributes. The validator.py module uses the schemas in the mwschema.py module for the validations that are easily handled by jsonschema. So, improvements in the mwschema.py module translate to improvements in the validator.py module. In addition, the validator.py module received substantial improvements outside of the improvements in the mwschema.py module. A host of consistency validations were added, for example validating that the subject sample factors in the SUBJECT_SAMPLE_FACTORS section and the METABOLITE_DATA section are the same. Many validations were added to detect duplicates, such as duplicate sample names, duplicate metabolite names, and duplicate rows in tables.

The most extensive validation, in terms of development time, was significantly improving and extending the validation on columns in the METABOLITES table. Previously, there was a validation that looked for column names that looked like certain standard column names and printed a message to the user to change it to the standard name. For example, “retention_time” would be a standard column name, but you can find variations on that name such as “ret time”, “ret_time”, or “retention time”. This validation was greatly expanded. Originally, 11 standard column names were validated, and that has now increased to 56. The method used to identify these column names was also improved, and an iterative test-improvement loop using all of the column names in the MW repository was done to fine tune the method for each name to reduce false positives and false negatives. Additionally, the validation was extended to also validate the values of each of these columns, not just the name. A similar test-improvement loop was used to fine tune the value validation. These extensive improvements spawned the new module, metadata_column_matching.py. This new module contains all of the new classes, functions, and resulting class objects used by validator.py to perform the standard column name and value validation. Since the code in the metadata_column_matching.py module could be used to validate the names and values of any kind of data in tabular form, we added extensive documentation (https://moseleybioinformaticslab.github.io/mwtab/metadata_column_matching.html) on how to use this module outside of the mwtab package for that purpose.

The validation messaging also received an overhaul. Internally the validation is done on the internal representation of the mwTab file, and validation messages reflected that in that the previous version. This caused the messages to make a bit less sense when referring to mwTab files in the tabular form and also caused a lot of repeating spurious messages about what was really 1 error. Now, validation is aware of whether it is validating a tabular or JSON version and crafts messages specific to the file type. The spurious repeating messages have also been condensed to just 1 message. The last noteworthy change to validation is that it no longer just prints messages. It now also returns a list of dictionaries, where each dictionary is a validation message along with tags categorizing the message, a unique ID, short name, and keys for the section and sub-section that generated the message. This change was largely motivated by the changes made to the mwFileStatusWebsite and made it much easier to improve the information provided by the website.

### 2.3. mwFileStatusWebsite Updates

Updates to the mwFileStatusWebsite were not quite as extensive as those to the mwtab package, but were required due to the extensive changes to the mwtab package. Since the website had to be updated due to the changes in the mwtab package, other maintenance or modernization updates were also done at the same time. These updates included switching the installation method to using a pyproject.toml file [13] instead of the older setup.py file, switching to a src distribution package organization, and switching to versioning using GitHub tags and the setuptools-scm package [14]. Some other minor updates were in the site functionality. Previously, when a user clicked on a specific study or analysis to see more detailed information, only one set of study or analysis details could be open at a time. We changed the cascading style sheets (CSS) so that any number of studies or analyses details can be displayed at the same time. The largest and most impactful changes came downstream from the changes to the mwtab package. Previously, when the mwtab package had many fewer validations, there were many more mwTab files that reported no validation issues and were put in the “passing” category. However, after adding many more validations to the mwtab package, most mwTab files have some sort of validation issue. In order to make the site a little more useful, we decided that it needed to include new categories and sub-categories of validation issues. To that end, we added a “Warnings Only” category which is for mwTab files that only have warnings in the validation output, and we added the sub-categories of “Value Error”, “Consistency Error”, and “Format Error” to sub-categorize the validation errors. A “Value Error” is where the value of a subsection or column is incorrect. For example, putting ‘5 W’ for a subsection asking for a voltage. A “Consistency Error” is where the same values should be in 2 different locations, but they are not. For example, when there are metabolites in the METABOLITES table that are not in the DATA table. A “Format Error” is where the file does not follow the mwTab specification. For example, the file has sub-sections that don’t exist or is missing required subsections. The desire to add these additional sub-categories to the website drove the changes to the mwtab package to support them, so the mwtab package changes drove changes to the website, and the website then drove changes to the mwtab package. Having the validation output in the previously described list of dictionaries was the best way to support these updates to the website. Previously, the website was parsing the text output from validation, but now it is much easier to utilize the list of dictionaries output to get the category and sub-category information needed by the website. Figure 1 illustrates the additional category and sub-categories, and you can visit the website at https://moseleybioinformaticslab.github.io/mwFileStatusWebsite/ to see its functionality. We also took the time to add unit testing to the site code, such that over 99% of the code lines are now covered.

### 2.4. Evaluation of the Metabolomics Workbench Repository

The mwtab package functionality was evaluated on every available mwTab and JSON formatted file available from MW (as of 22 October 2025). A total of 6125 analysis IDs were available, and the downloading of both mwTab and JSON formatted data files was attempted through the MW REST interface.

The files from MW were evaluated for a number of formatting, value, and consistency errors/issues. First, the successful download of each analysis file through MW’s REST API was tested in both mwTab and JSON format. Second, the successful parsing of each downloaded file into MWTabFile objects was tested. Third, each parsed file was validated using the validation methods in the mwtab Python package. Fourth, the consistency between parsed data from mwTab and JSON formats was tested.

Downloading was done only using the new version of the mwtab package, but parsing and validation steps were performed using both the previous version of the mwtab package and the new version. Consistency between the mwTab and JSON formats parsed by both the new and previous versions of the mwtab package were tested using the updated consistency checking code in the mwFileStatusWebsite.

## 3. Results

### 3.1. Evaluation of the Metabolomics Workbench Repository

As of 22 October 2025, a total of 6125 analyses were available for download through MW’s REST interface. With the mwtab 2.0.0 version, we applied a series of evaluations on each analysis in both mwTab and JSON formats. These evaluations were performed in a natural progression starting with whether the analysis file could be downloaded and parsed, followed by whether the mwTab and JSON files were consistent with each other, and finally applying an extensive list of validation tests.

#### 3.1.1. Analysis IDs with Files Missing from the Metabolomics Workbench

When we attempted to download all available analyses, a number of analyses were not present for download in a given format. Of those analyses, 8 could not be downloaded in mwTab format and 6 could not be downloaded in JSON format. The specific analysis IDs are given in Table 2. Only blank pages were present for these files.

**Table 2.** List of analysis IDs and the format of files that could not be downloaded through Metabolomics Workbench’s REST API on October 22, 2025.

| File Format | Analysis ID |
| --- | --- |
| mwTab | AN002312, AN005082, AN005098, AN005099, AN005557, AN006051, AN006148, AN006593 |
| JSON | AN005098, AN005099, AN005557, AN006051, AN006148, AN006593 |

It can be seen by comparing the analysis IDs in Table 2 that 6 out of the 8 files are shared between both formats. Further investigation was done by finding these analysis IDs on the Metabolomics Workbench website through the summary of all studies. The files available for download that way were not all blank like they are when using the REST interface, but when they are not blank, they are also not in the correct format. For example, the mwTab format file for AN002312 contains only the tables that would be in the DATA and METABOLITES sections. It is not clear why most of the analysis IDs that could not be downloaded overlap across formats but not all of them. This seems to imply a lower-level issue specific to each analysis ID, since most are not available in either format.

#### 3.1.2. Analysis Files Which Could Not Be Parsed

Of the 6117 downloaded analyses in mwTab format, 51 files could not be parsed into MWTabFile objects. Of the 6119 downloaded analyses in JSON format, 124 files could not be parsed into MWTabFile objects. For 37 analysis IDs, both the mwTab formatted file and the JSON formatted file were not able to be parsed.

Of the 51 mwTab files which could not be parsed, 30 files contained formatting errors in the additional data section of the SUBJECT_SAMPLE_FACTORS where either the semicolon delimiter between key value pairs was missing, or there was an additional ‘=’ character that should be separating the key and value. 13 files had formatting errors where there were lines without the capitalized sections at the beginning, this was usually due to the value of a section containing a value with a carriage return. 5 files had formatting errors where they had a line that was missing the tab after the capitalized section at the beginning of the line. 2 files had formatting errors where the factors had an additional ‘:’ character that should be separating the key and value. 1 file had a very badly formatted METABOLITES_DATA table that had multiple MS_METABOLITE_DATA_END lines between the rows of the table. There should only be one MS_METABOLITE_DATA_END line at the very end of the table.

Of the 124 JSON files which could not be parsed, 69 files were simply badly formatted JSON. For instance, if the mwTab formatted version of the file had an extra ‘=’ character as described above, then that would translate into an extra invalid character in the JSON version. 48 files had non-dictionary values in the tabular data. The METABOLITES_DATA and METABOLITES tables within the mwTab formatted version get translated into a list of dictionaries in the JSON version, but for some files one or more of these dictionaries instead have a non-dictionary value like “false”. When compared to the mwTab formatted version of the same file, the non-dictionary values correspond to a row in a table and the value for that row is not “false”. This means that the JSON version is missing data that is in the mwTab version. We believe that the JSON files are created by parsing the mwTab version of the file and that for some reason certain rows fail to parse correctly and result in the “false” value. By default, the mwtab package will not parse these files, but there is now in version 2.0.0 a “force” CLI option and API function parameter that will instead ignore these values and allow the files to be parsed. The last 7 files that could not be parsed are due to having duplicate keys at the highest level of the JSON. For instance, having 2 “MS” or “MS_METABOLITE_DATA” keys. Although one of the improvements in this release is being able to handle duplicate keys, that is only for values in lower levels of the data, such as Factors within the SUBJECT_SAMPLE_FACTORS. Duplicate keys at the highest level are much harder to deal with. Also, if you look at the mwTab formatted version of these 7 files, the duplicated keys in the JSON do not correspond to duplicate sections in the mwTab.

The mwtab package has a “convert” command that allows a user to convert between the mwTab formatted version of a file and the JSON version of a file and vice versa. When we use this command to convert all of the downloaded mwTab formatted files into JSON files, all of the converted JSON files are parsable. In other words, converting the mwTab formatted files to JSON files fixes 100% of the JSON file parsing errors for the files that have a mwTab formatted version without parsing errors.

For a full list of files with parsing errors, see Supplemental Table S1.

#### 3.1.3. Comparing Parsability Between mwtab Version 1.2.5 and Version 2.0.0

When using the previous version of the mwtab package (1.2.5), 162 analysis files in the mwTab format could not be parsed into MWTabFile objects, and 69 analysis files in the JSON format could not be parsed into MWTabFile objects.

Of the 162 files in the mwTab format, 20 overlap with the 51 unparsable files when using the new 2.0.0 version. The 31 files that are parsable when using the previous version of the package have arguably a worse issue. Although the package will parse them, the resulting data will be wrong and won’t match the file that was read in if the package was used to write out what it had parsed. The root cause is the aforementioned issue in the SUBJECT_SAMPLE_FACTORS where some of the syntax characters can be missing or there are extra ones due to the data uploader using those characters in the key or value. The previous version of the mwtab package used a more naïve approach to parsing the SUBJECT_SAMPLE_FACTORS that was unaware of this issue, resulting in incorrect MWTabFile object representations. Naïve approaches to parsing explain the other 142 unparsable files. Detailing every naïve assumption that causes an issue in parsing is not within the scope of this paper, but 112 come from the SUBJECT_SAMPLE_FACTORS and the other 30 come from tab placement issues throughout the sections.

Of the 69 files in the JSON format, all of them overlap with the 124 unparsable files when using the new 2.0.0 version. The 55 files that are parsable by the previous version and not the new version are the 48 files with non-dictionary table values and the 7 files that have duplicate keys at the top level. Even though the new version of the package results in less files being parsable, it is better overall because those issues in a file can cause many more pernicious issues deeper in a data workflow if they were parsed in with the more naïve method used in the previous version of the package. Also, 48 of the 55 files can be parsed in if the aforementioned “force” option is utilized.

We will also mention here that 10 of the files in the mwTab format and 24 of the files in the JSON format cannot be validated when using the previous version of the package because the package will crash. This is partly due to naïve coding assumptions and partly due to files from the Metabolomics Workbench not being as reliably formatted as you might like or expect. The issues causing the validation code to crash are due to certain assumptions that would be reasonable and would be expected of many data providers. For example, in a JSON object, data that is not present can be represented by having the key with a null value, or by not including the key. A reasonable expectation is that a data provider would choose one of those methods and then use that as the standard across their datasets. They would not have some data sets that had keys with null values and some without the key. Unfortunately, data sets from the Metabolomics Workbench do have this issue. The new version of the mwtab package fixed all of these errors in the validation code and does not crash on any of the 6117 files in the mwTab format or any of the 6119 files in the JSON format.

For a full list of files with parsing errors and validation crashes see Supplemental Table S2 and Supplemental Table S3.

#### 3.1.4. Consistency Errors Between mwTab and JSON Formatted Files

We compared the internal package representations when reading in the downloaded mwTab version of the analysis with the downloaded JSON version of the analysis to see how consistent they are between each other. In this context, “consistent” means that the

MWTabFile objects contain the same data. We did this comparison for both the new 2.0.0 version of the mwtab package and the previous 1.2.5 version. For the new version, of the 5979 parsable, non-missing files, 3077 were consistent with each other and 2902 were inconsistent with each other. For the 1.2.5 version, of the 5895 parsable, non-missing files, 2944 were consistent with each other and 2951 were inconsistent with each other. The reasons for the inconsistencies were investigated. Figures 2 and 3 show a summary of the categorized reasons for inconsistency for the new and previous versions of the mwtab package, respectively.

**Figure 2.**
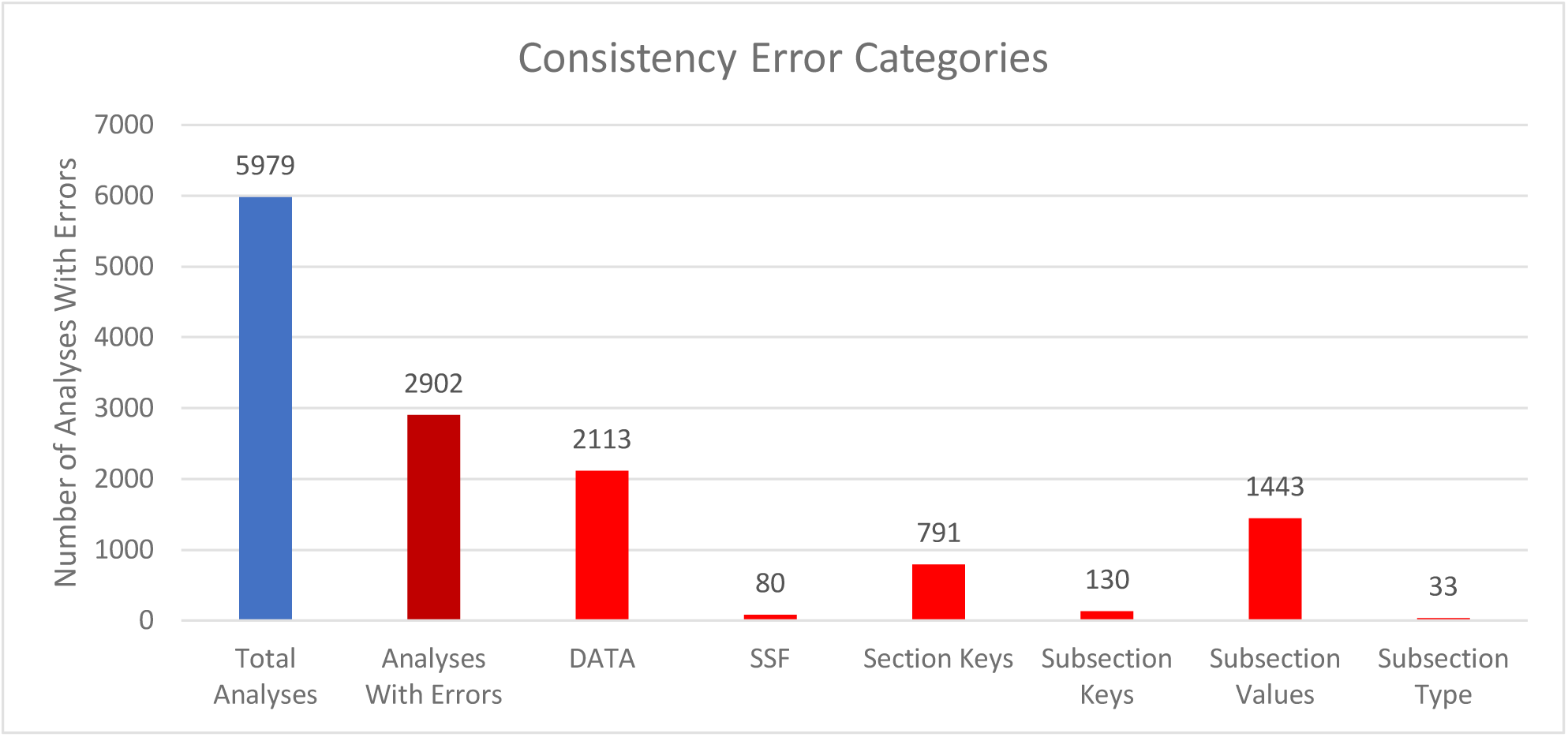
Column chart showing the breakdown of reasons for the inconsistencies between downloaded mwTab formatted and JSON formatted files for the new 2.0.0 version of the mwtab package.

**Figure 3.**
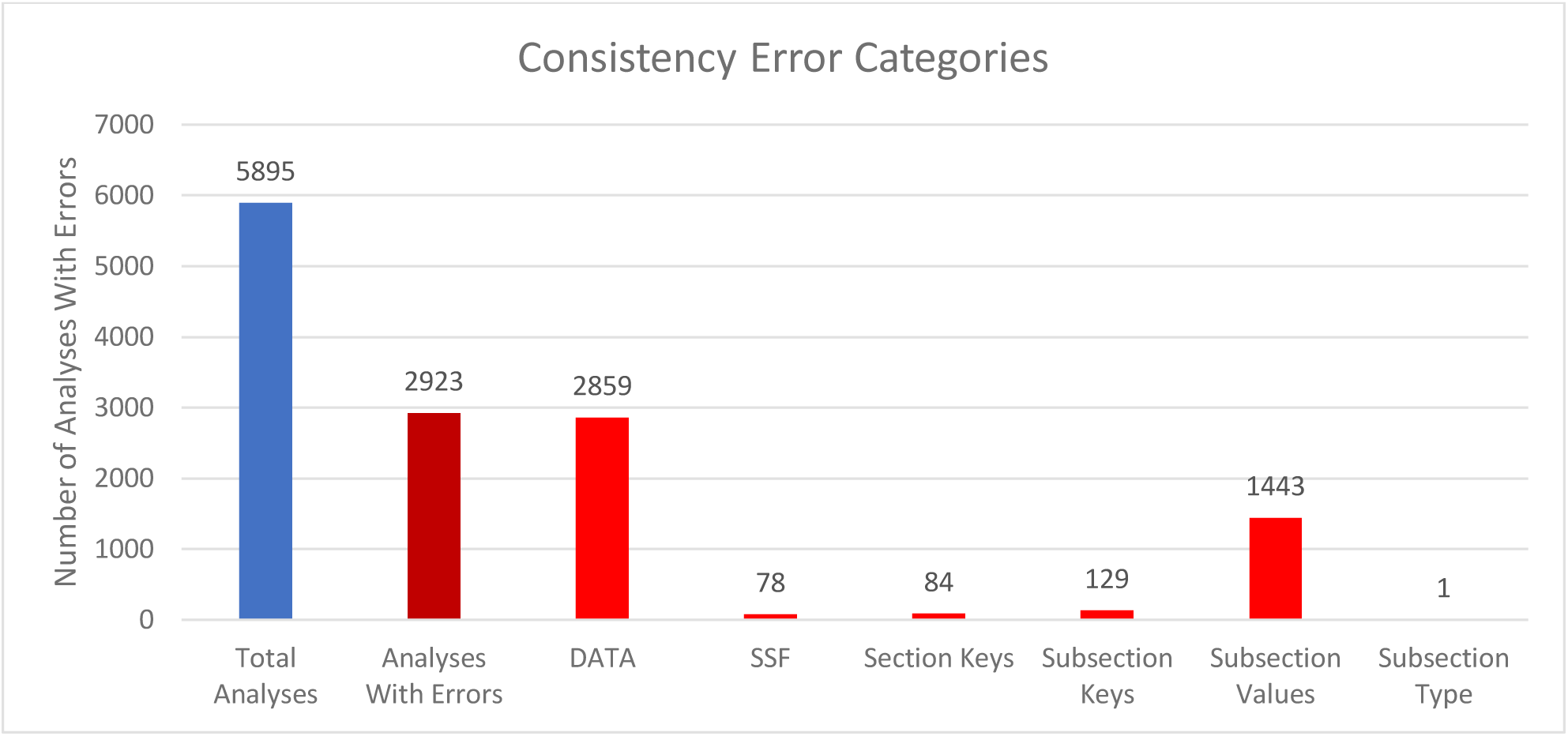
Column chart showing the breakdown of reasons for the inconsistencies between downloaded mwTab formatted and JSON formatted files for the previous 1.2.5 version of the mwtab package.

The “DATA” category refers to the 2 formats having differences in the DATA or METABOLITES sections. Similarly, “SSF” refers to the 2 formats having differences in the SUBJECT_SAMPLE_FACTORS section. “Section Keys” means there were differences between the sections, for example one may have a CHROMATOGRAPHY section and one may not. Similarly, “Subsection Keys” means there were differences within a section, for example one might have a RESULTS_FILE key in the MS section and one may not. “Subsection Values” means that for the same subsection within a section the values were not the same between the 2 formats. Lastly, “Subsection Type” is similar to “Subsection Values”, but not only is the value different, the data type is different. For example, a subsection might have a string (text) value in one format, but a list of strings in the other.

Comparing Figures 2 and 3 closely, there are 2 significant discrepancies between the new and previous versions of the mwtab package. The new version has significantly more “Section Keys” inconsistencies and the previous version has significantly more “DATA” inconsistencies. Nearly all of the extra “Section Keys” inconsistencies come from 736 mwTab formatted files that have a “#FACTORS” line in them. There is not supposed to be a FACTORS section at all, but for some reason these files have a line like that in them. This causes the mwtab package to create an empty FACTORS section in the MWTabFile object when reading in that file, but the corresponding downloaded JSON version of the file does not have that section, thus the inconsistency. These don’t show up in the previous version of the mwtab package because of a small difference in parsing. The previous version does not create the FACTORS section, because it is empty, but this behavior can cause other issues, so it was changed in the new version. The extra “DATA” inconsistencies also come from a difference in parsing. The mwTab formatted versions have tab delimited data tables for the DATA and METABOLITES sections. There are 2 different kinds of tab issues that can happen with these tables. One is that there can be extra tabs with no data tacked on to the end of rows, and not necessarily every row, such that the table can look like it has several extra columns with no name and no values. The other tab issue that can happen is that when a row has a blank value for the last column(s), the tabs that are supposed to be there are not there. This makes it look like the row only has some of the columns. The previous version of the mwtab package parsed these tables more naively and relied on the number of tabs in each row being correct. The new version of the package is not so naïve and harmonizes the columns for each row, making it more consistent with the downloaded JSON.

We also used the mwtab package’s “convert” command through its CLI to convert the downloaded mwTab formatted files into JSON formatted files and compared them. Unsurprisingly, they are 100% consistent. This is another reason why we recommend working with our converted JSON files over ones directly downloaded.

The mwFileStatusWebsite does this consistency check between the downloaded mwTab and JSON formatted files every week along with its validation of the files, and was updated to fix some inaccuracies and address changes made in the new version of the mwtab package.

#### 3.1.4. Validation Issues

The improvements in validation are the most extensive changes made in the new version of the mwtab package. Table 3 shows the number of analyses that fall into a few general validation issue categories when using the previous version of the mwtab package to validate.

**Table 3.** The number of analyses that fall into each category for each format when using the previous 1.2.5 version of the mwtab package to validate the analyses.

| Issue Category | mwTab | JSON |
| --- | --- | --- |
| Passing | 5497 | 4907 |
| Missing | 8 | 6 |
| Parsing Error | 162 | 69 |
| Validation Bug | 10 | 24 |
| Errors | 456 | 1125 |

Table 4 shows a similar table to Table 3 but expanded into more categories when using the new version of the mwtab package, and Table 5 gives more in-depth descriptions for each category.

**Table 4.**
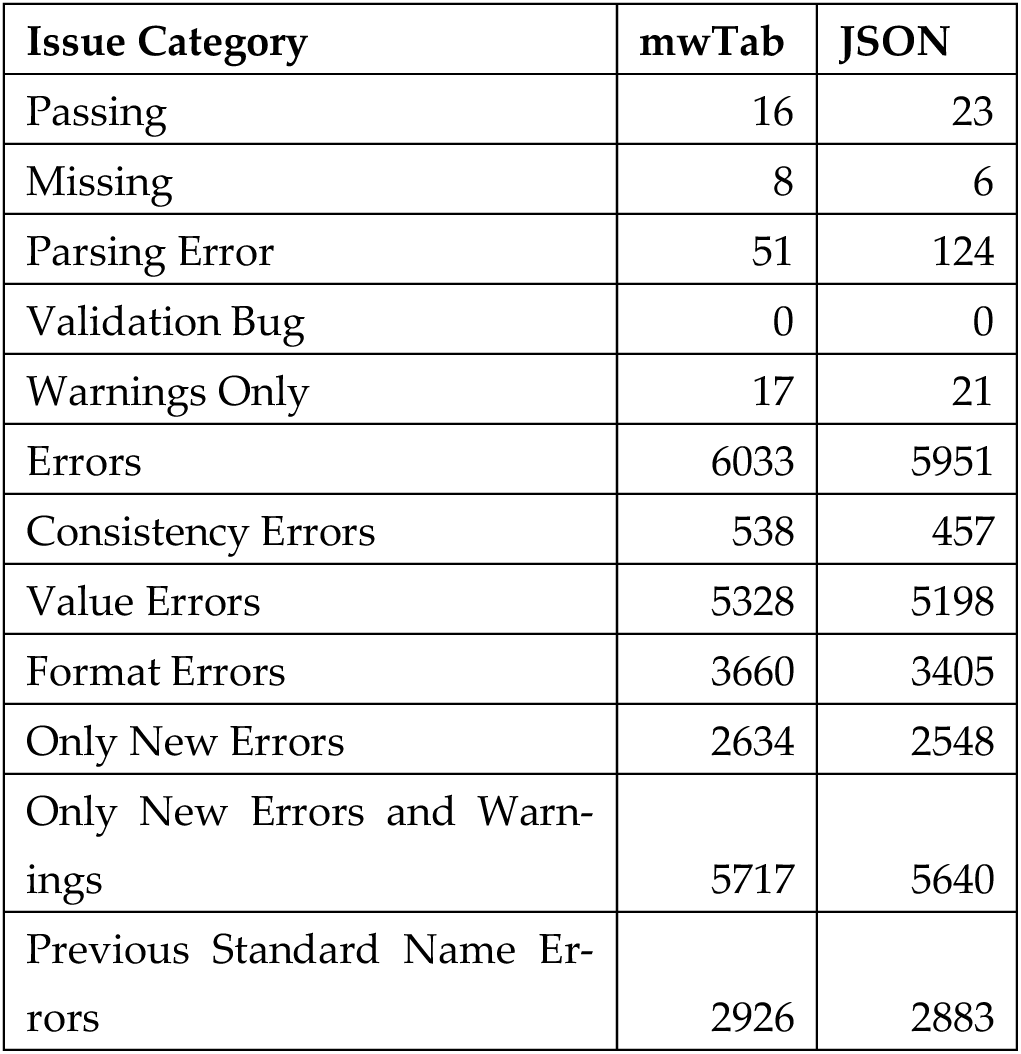
The number of analyses that fall into each category for each format when using the new 2.0.0 version of the mwtab package to validate the analyses.

| Issue Category | mwTab | JSON |
| --- | --- | --- |
| Passing | 16 | 23 |
| Missing | 8 | 6 |
| Parsing Error | 51 | 124 |
| Validation Bug | 0 | 0 |
| Warnings Only | 17 | 21 |
| Errors | 6033 | 5951 |
| Consistency Errors | 538 | 457 |
| Value Errors | 5328 | 5198 |
| Format Errors | 3660 | 3405 |
| Only New Errors | 2634 | 2548 |
| Only New Errors and Warnings | 5717 | 5640 |
| Previous Standard Name Errors | 2926 | 2883 |

**Table 5.** Description of each validation issue category given in. Tables 3 and 4.

| Issue Category | Description |
| --- | --- |
| Parsing Error | File could not be parsed. |
| Validation Bug | A coding error was encountered during validation. The file did not necessarily have something wrong with it. |
| Passing | File had no validation issues. |
| Missing | File could not be downloaded. |
| Warnings Only | All of the validation issues were only warnings. |
| Errors | There was at least 1 validation issue with the file that was not a warning. |
| Consistency Errors | There was at least 1 validation issue due to inconsistencies between data in the file. |
| Value Errors | There was at least 1 validation issue due to incorrect value(s) in the file. |
| Format Errors | There was at least 1 validation issue due to the file not meeting the mwTab specification. |
| Only New Errors | Every issue found is one that has been added in this latest version of the mwtab package. |
| Only New Errors and Warnings | Every issue found is one that has been added in this latest version of the mwtab package or is a warning. |
| Previous Standard Name Errors | If the previous version of the mwtab package had validated while using its code for standard name matching, then the file would have errored instead of passing. |

The categories had to be expanded in Table 4 for the new version of the mwtab package, because “warnings” are a new concept in this version, as are the “consistency”, “value”, and “format” categories of validation issues. “Warnings Only” was added as a new category in the mwFileStatusWebsite, since many new validation warnings were introduced in the new version of the mwtab package. Warnings are less severe than an outright error and have a chance of being a false positive. We felt adding this category would add more gradations and nuance when evaluating files from MW. We also added categories to the validation issues, “consistency”, “value”, and “format”. The “consistency” category in this context refers to consistency between sections within the file. For example, if there is a Sample ID in the METABOLITE_DATA table that is not in the SUBJECT_SAMPLE_FACTORS section, then there is a consistency error between those 2 sections. A “value” error is when there is an issue with the value somewhere. For example, if there are 2 metabolites in the METABOLITES table with the same name, then that is a value error because one of the values is likely incorrect. Lastly, a “format” error is when there is an issue that violates the mwTab specification. For example, if the required sub-section SUBJECT_TYPE is not present in the SUBJECT section, that is a format error because the mwTab specification requires this field to be present. The prior versions of the mwtab package had primarily focused only on “format” errors.

The “Only New Errors”, “Only New Errors and Warnings”, and “Previous Standard Name Errors” categories were added to Table 4 to help show how so many files went from “Passing” in the previous version of the mwtab package, to “Errors” in the new version. We can see from “Only New Errors” that around 2600 files are now not passing solely because of new validations introduced. We can then see from “Only New Errors and Warnings” that around 5700 files are now not passing due to new validations or warnings. The gap between “Only New Errors” and “Only New Errors and Warnings” is explained with the “Previous Standard Name Errors” category, but some preamble is required to understand this category. The previous version of the mwtab package had a less sophisticated version of a validation that was greatly expanded in the new version of the package. This validation looks for certain standard columns in the METABOLITES table that do not have the corresponding standard name and prints a warning to the user to rename it. For example, one standard column name is “moverz_quant”, but this could appear in data sets as “mz”, “m/z”, “moverz”, etc. In the new version of the mwtab package, this validation is much more robust for the names that were covered in the previous version and covers many more names. This validation in the new version also always prints a warning to the user, but in the previous version it was an optional validation and it was printed as an error instead of a warning. The results in Table 3 were determined with this validation turned off, so the “Previous Standard Name Errors” category in Table 4 is the number of analyses that would have been in the “Errors” category in Table 3 if the validation had been turned on. Since this validation is technically not new, files that fail the validation would not appear in the count for the “Only New Errors” category but do get counted if we also add in warnings. The reason the count for the “Only New Errors and Warnings” category does not exactly match the count for the “Passing” category in Table

3 is because there are some other smaller issues that account for the difference. For example, in the previous version of the mwtab package blank Subject IDs in the SUBJECT_SAMPLE_FACTORS would be a validation error, but in the mwTab specification Subject IDs are optional, so they are no longer errors in the new version of the package. Issues like this affect far fewer data sets, so they were not extensively tabulated and accounted for in Table 4. The majority of data sets that went from Passing to Errors did so due to new validations and making the standard name validation mandatory.

Table 6 shows the number of analyses that failed each validation ID, and Table 7 gives a description of each validation.

**Table 6.**
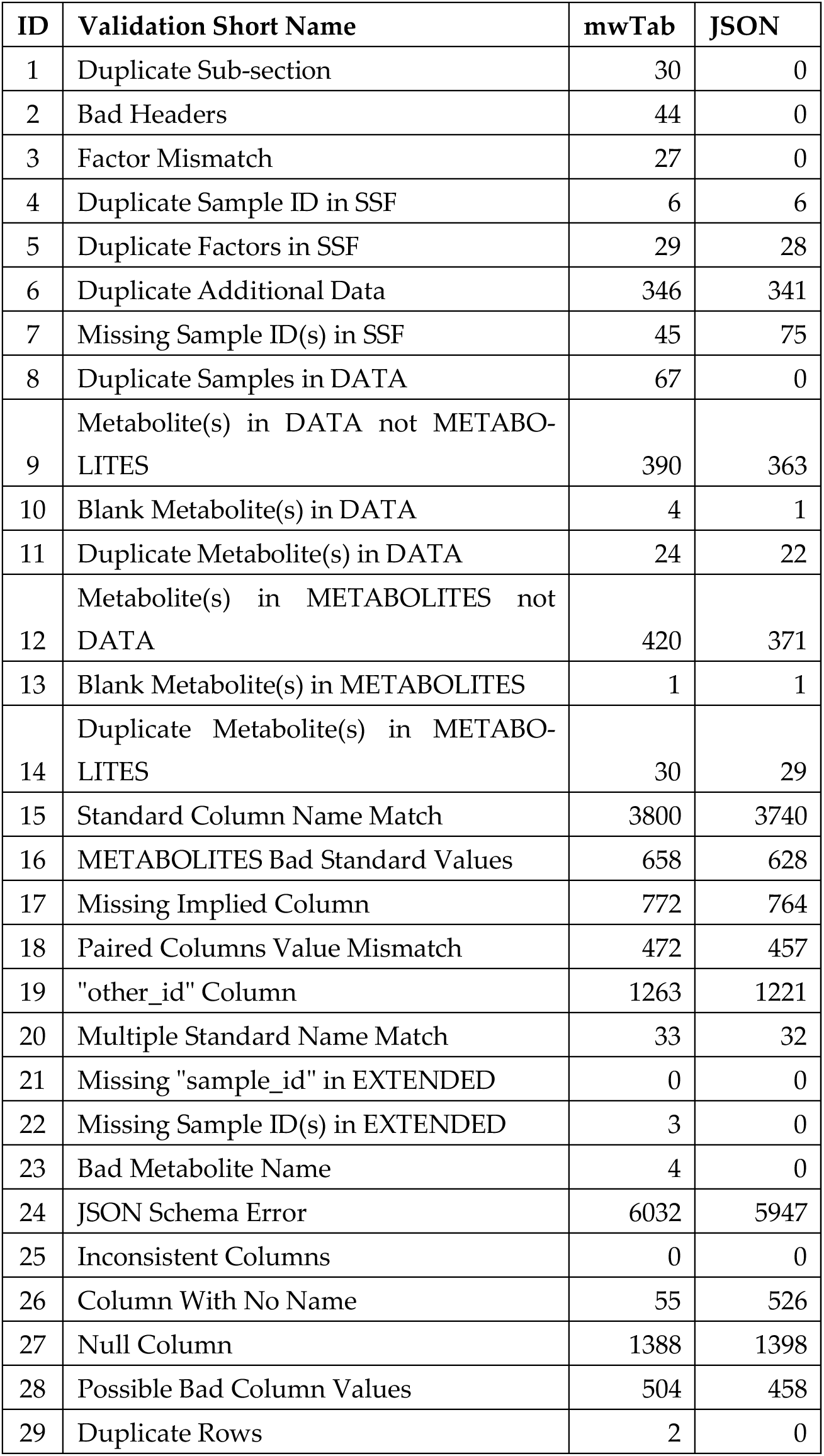
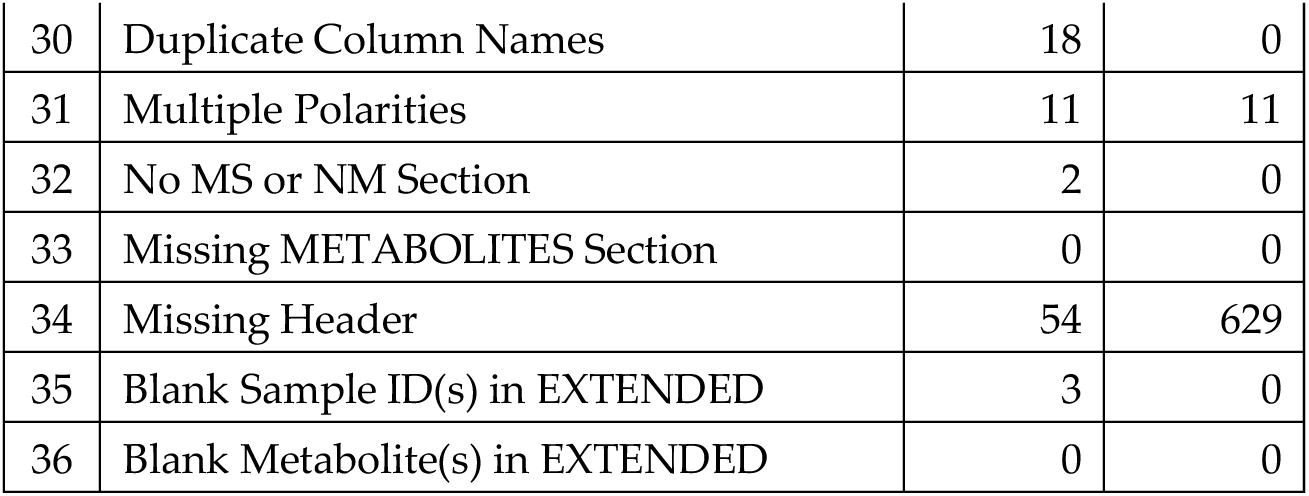
The number of analyses that failed each validation for both downloaded formats.

**Table 7.**
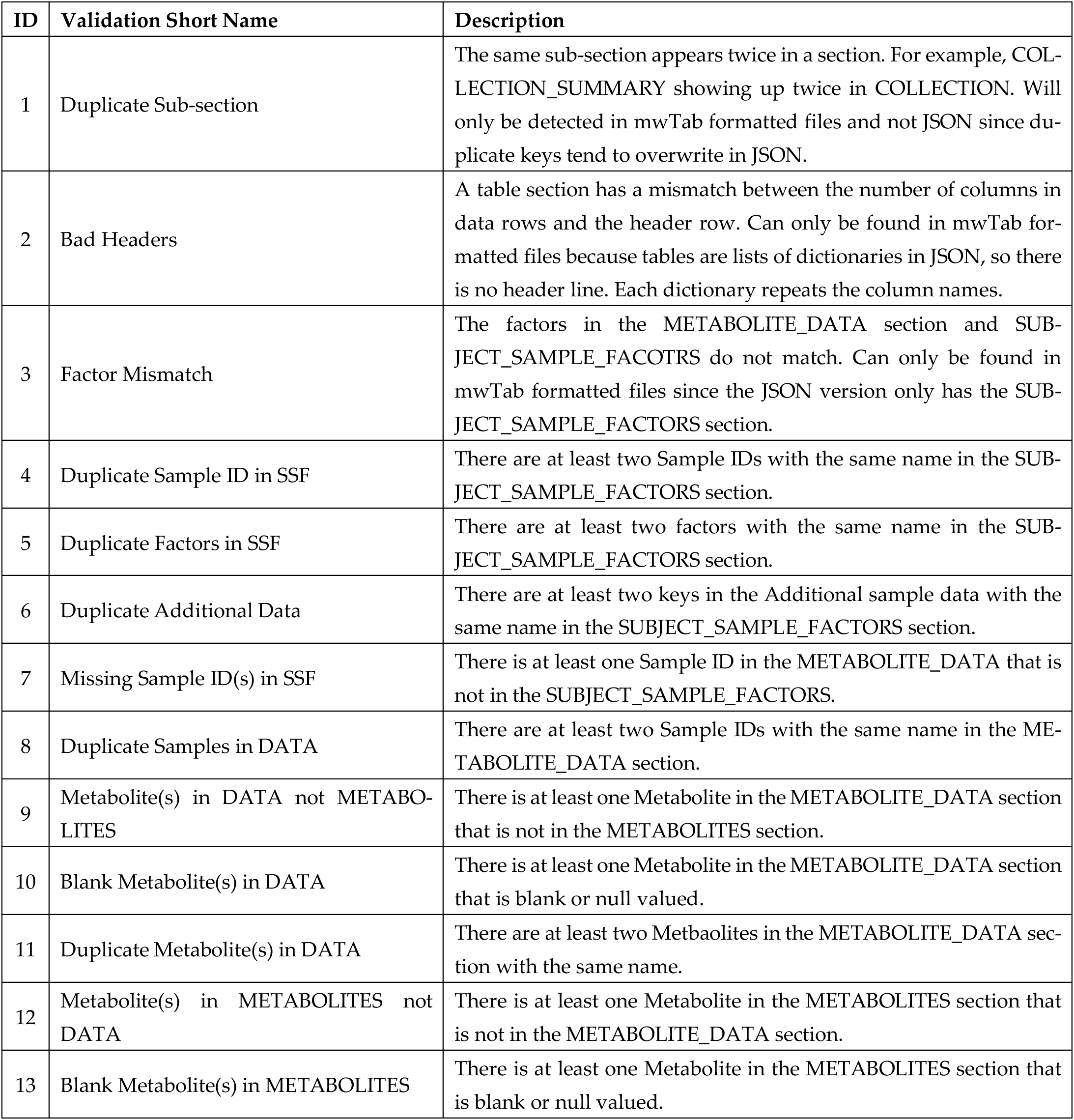
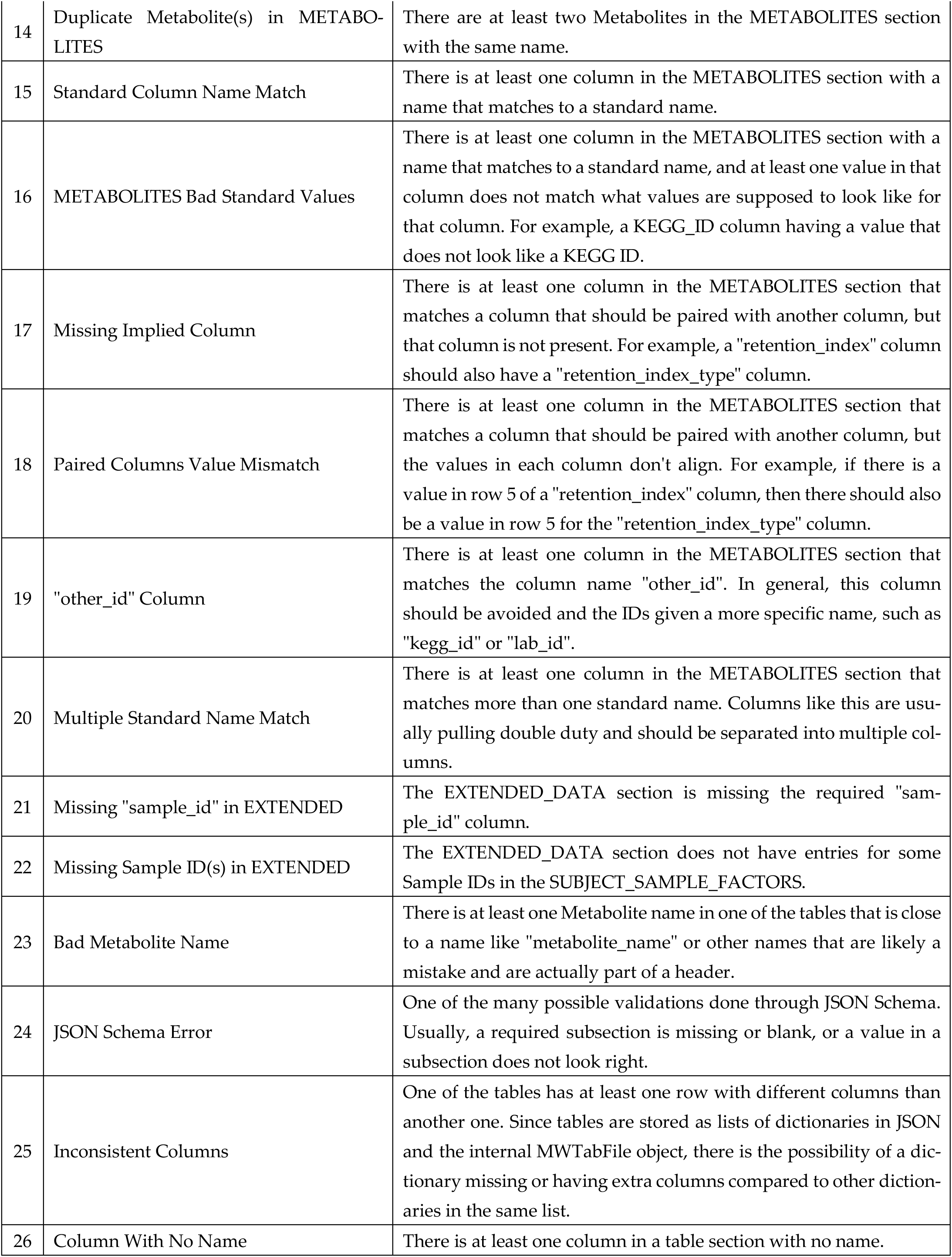
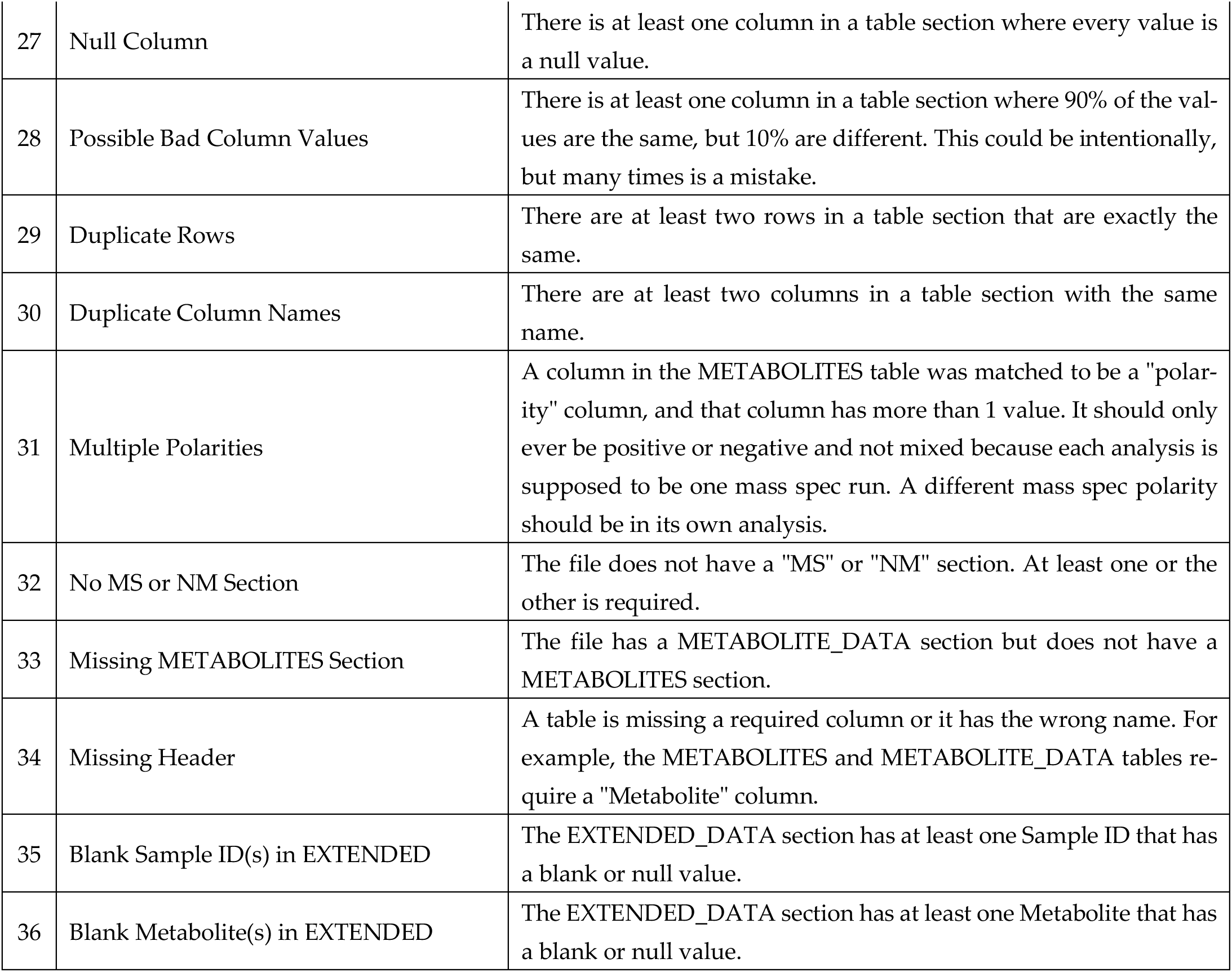
Descriptions of the validations.

The number of analyses that fail each validation varies wildly, from 0 failing to almost all failing. The validations with 0 analyses failing can usually be explained by implication to another validation. For example, take a validation, say validation B, that is similar to another validation, say validation A, and validation A was added because it was seen in actual datasets, but validation B was added because of its similarity to validation A. Since validation A was confirmed to be an issue, it implies that validation B could also be an issue. For example, “Blank Metabolite(s) in EXTENDED” was added because this issue was found in the METABOLITES and DATA tables but was not directly found in the EXTENDED table. The validations with much larger numbers of analyses failing are “Standard Column Name Match” and “JSON Schema Error”. “Standard Column Name Match” is explained by there just genuinely being that many data sets with a METABOLITES table that has at least one column with a non-standard name. “JSON Schema Error” has such a large number, because it represents many validations under one label. The mwtab package uses the jsonschema package to do some validations and due to how jsonschema works within mwtab it is easier to assign one ID to those validations than it is to try and assign an ID to validations within jsonschema. An analysis of the JSON Schema validations showed that the biggest sources of failed validations came from incorrect subsection values, null valued sub-section values, and missing required sub-sections, in that order.

Careful observation of Table 6 will also lead to seeing some large discrepancies between the number of analyses failing between the mwTab and JSON formats. The top 3, “Duplicate Sub-section”, “Bad Headers”, and “Factor Mismatch” are the easiest to explain. They are issues that by their very nature can only occur in the mwTab format or at least are much less likely to occur in the JSON format. A duplicate sub-section would be when a section, say SUBJECT, has a sub-section repeated, for example SUBJECT_TYPE. The repetition could have the same information or different information. This is much less likely to occur within the JSON format because sub-sections are keys in a JSON object and typically if a key with the same name is added to a JSON object, instead of duplicating it will just overwrite the value for that key. “Bad Headers” occur when there are different numbers of columns and data in rows within a table section. For example, the table in DATA might have 5 columns in the header row, but then a data row might have 7 pieces of data. This doesn’t happen in the JSON format because those tables are stored as lists of JSON objects and each object always has a 1 to 1 relationship with columns and data. A “Factor Mismatch” occurs when the factors in the SUBJECT_SAMPLE_FACTORS section and the factors in the DATA section are not the same. This cannot occur in the JSON format because it only has the SUBJECT_SAMPLE_FACTORS section, there is no second source of factor information that can be mismatched.

The next discrepancy is “Duplicate Samples in DATA”. This occurs when one or more sample names are used twice as column names in the DATA section. The reason that this does not occur in the JSON format is similar to the explanation already given for the “Duplicate Sub-section” validation. The JSON format does not duplicate the sample keys in its objects used to store the table data, it is instead overwritten so that only the last sample with that name will have its data in the JSON format. This means that information is lost when using the downloaded JSON. This also differs from how MW used to construct the JSON in this situation. We have JSON downloads from April 2024 for these datasets where the key was duplicated in the JSON file. These duplications led to one of the more significant improvements in the mwtab package to allow it to read and write these duplicate keys without losing data. When using the mwtab package, you can read and write datasets that have this duplicate key issue without any problems, and JSON files with this issue that are written out using the mwtab package do not lose data. This is another reason why we do not recommend working with JSON directly downloaded from MW and instead recommend converting mwTab to JSON using the mwtab package.

The next two discrepancies are larger in magnitude, and this time affect the JSON format more than the mwTab format. “Column With No Name” is simply when one of the table sections has a column with no name. Typically, this column is the very last column and is not just a column with no name, but is also a column with all null values. The mwTab format does not have these columns because during parsing empty values at the end of lines are removed so the columns don’t end up in the internal representation. The discrepancy in this validation does not translate into a discrepancy in the “Null Column” validation, because so many of these datasets with an extra null column at the end also have other null columns. A “Missing Header” occurs when a table is missing one of its required headers, typically “Metabolite”. This occurs more in the downloaded JSON, because for whatever reason there are several data sets that have “metabolite_name” instead of “Metabolite”. The mwTab version of the file doesn’t have the issue because during parsing the table information is always built using the correct name and does not use whatever is in the file. The JSON version is simply read in as is, however. Both of these discrepancies go away if using JSON files converted from the mwTab formatted files using the mwtab package.

Table 8 shows the number of analyses that have a validation issue for each section for each format.

**Table 8.**
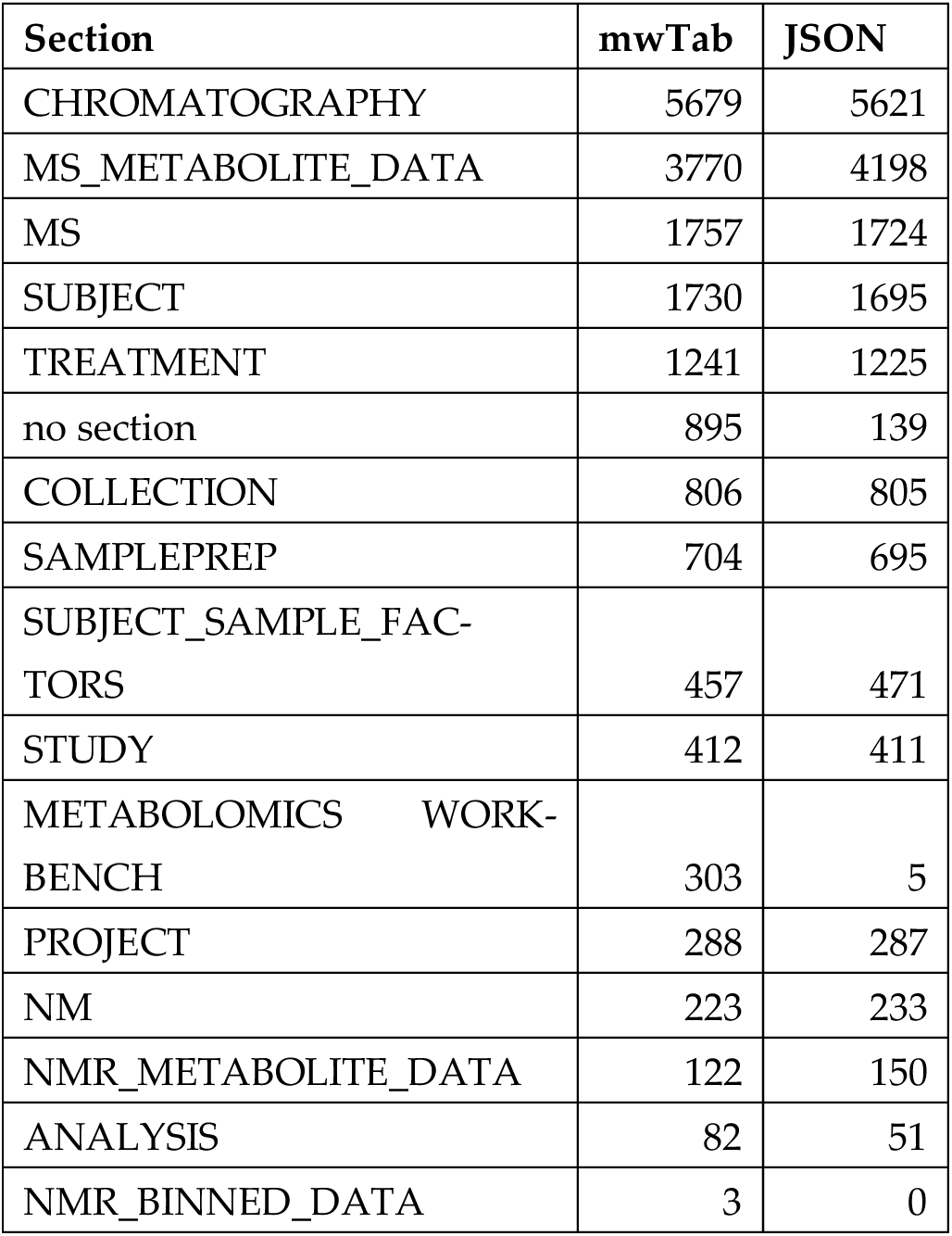
The number of analyses that have a validation issue for each section for both downloaded formats, sorted from largest to smallest.

| Section | mwTab | JSON |
| --- | --- | --- |
| CHROMATOGRAPHY | 5679 | 5621 |
| MS_METABOLITE_DATA | 3770 | 4198 |
| MS | 1757 | 1724 |
| SUBJECT | 1730 | 1695 |
| TREATMENT | 1241 | 1225 |
| no section | 895 | 139 |
| COLLECTION | 806 | 805 |
| SAMPLEPREP | 704 | 695 |
| SUBJECT_SAMPLE_FACTORS | 457 | 471 |
| STUDY | 412 | 411 |
| METABOLOMICS WORKBENCH | 303 | 5 |
| PROJECT | 288 | 287 |
| NM | 223 | 233 |
| NMR_METABOLITE_DATA | 122 | 150 |
| ANALYSIS | 82 | 51 |
| NMR_BINNED_DATA | 3 | 0 |

The section with the greatest number of analyses failing is CHROMATOGRAPHY, but this is somewhat misleading because an update to the specification added more required fields to this section after several thousand datasets had already been uploaded without those fields. The second highest on the list is not surprising since MS_METABOLITE_DATA encompasses both the DATA and METABOLITES tables and many of the new validations focus on validating the tables in various ways. Within this section the validations that were failed the most were “Standard Column Name Match”, “Null Column”, and “Possible Bad Column Values”. The two major discrepancies between the formats in this table is in the “no section” category and METABOLOMICS WORKBENCH section. The reason for the discrepancy in “no section” is due to many mwTab formatted files having a #FACTORS line in them. This is not a valid section name and should not be in any file for any reason. The corresponding JSON versions of the file do not have the empty FACTORS section. The reason for the discrepancy in the METABOLOMICS WORKBENCH section is that for whatever reason the PROJECT_ID subsection is blank for many of the mwTab formatted files, but not for their corresponding JSON versions.

#### 3.1.6. Conversions and Roundtripping

We have mentioned several times that we used the mwtab package to convert the mwTab formatted files into JSON formatted files and highlighted how those are often superior to the directly downloaded files, but in our testing of the package we also did other conversions. A “roundtrip” in this context refers to reading in a file, writing it back out as one of the 2 formats, and then reading it back in with no loss of information. We did this roundtrip testing from mwTab to mwTab, mwTab to JSON, JSON to mwTab, and JSON to JSON. All conversions were able to be written out and read back in without errors or loss of information except for 2 files in the mwTab to mwTab conversion. AN004560 and AN005763. They both have the same issue. A #FACTORS line that should not exist. It causes the NMR_RESULTS_FILE line to be pulled into a FACTORS key that shouldn’t exist and then when it is written out the NMR_RESULTS_FILE line is slightly different and missing the two-letter code and semicolon (NM:) because there is no two-letter code for FACTORS. Not having this two-letter code then causes a parsing error when trying to read in the converted file. It is very difficult to parse in a way to avoid this error without coding for it specifically. We have instead decided to develop a new package, rcha_metab, for which the purpose is to repair, clean, harmonize, and augment (rcha) metabolomics files. This delegates the task of removing bad lines like the #FACTORS line, among other things, so that the mwtab package does not have to worry about those kinds of issues. This package is still in active development and not yet publicly available, but we hope to make it available soon.

## 4. Discussion

The early focus of the mwtab package was to improve programmatic and FAIR access of metabolomics data in MW and to facilitate creation of clean mwTab formatted files for deposition into Metabolomics Workbench. After working on large-scale reuse of MW datasets [15], it was clear that the mwtab package needed to be updated to facilitate this new use-case. Mainly, a wider range of validations are needed to identify issues that prevent or hinder large-scale reuse. Also, the new use-case spawned our development of another package, rcha_metab, to repair, clean, harmonize, and augment (rcha) the datasets for large-scale reuse [15]. Development of rcha_metab directly fed into the improvements made to version 2.0.0 of mwtab. Also, parsing was improved so fewer files failed to be read in. Error handling was improved so that one “bad” file did not interrupt the batch processing of multiple files. Also, the package is now able to handle duplicate keys so information is not lost. The validation facilities were greatly expanded and improved. Several validation categories were created and many more validation tests were added. Now, validation messages are more descriptive while spurious messages were reduced. The documentation was updated for all of these changes and can be found at https://moseleybioinformaticslab.github.io/mwtab/. Automated unit and integration testing was also improved and over 95% of code lines are now covered. All of these improvements also necessitated improving the mwFileStatusWebsite which runs the validations discussed here once a week on all downloadable analysis files in MW and then displays the results through its website.

Improvements in parsing in version 2.0.0 of mwtab reduced non-parsability from 162 mwTab formatted files to 51 mwTab formatted files while non-parsability of JSON formatted files increased from 69 to 124. However, the 55 JSON files that are parsable by the previous version and not the new version are the 48 files with non-dictionary table values and the 7 files that have duplicate keys at the top level, which causes significant issues later in internal representation and validation. Given the improvements in parsability of mwTab formatted files, the tradeoff is worth it, since the mwTab formatted files are more reliable for reuse.

With the large addition of validation tests in the 2.0.0 version of mwtab, far fewer analysis files are considered clean (i.e, without issues). Of the files that could be downloaded and parsed all but 16 mwTab formatted files and 23 JSON formatted files had some kind of validation issue, and all but 33 mwTab formatted files and 44 JSON formatted files had a serious validation error. This is in stark contrast with the previous version of the mwtab package where most of the mwTab formatted files (5497 out of 6117) had no validation issues and most of the JSON formatted files (4907 out of 6119) had no validation issues. We also showed a more in-depth breakdown of the largest sources of validation issues in terms of the precise validation IDs and the sections in the files. The validation ID with the most failing analyses was ID 24, JSON Schema Error, and the largest sources of failing analyses within that ID were validations on sub-section values and missing required sub-sections. The top 2 sections causing analyses to fail validation were CHROMATOGRAPHY and MS_METABOLITE_DATA. Although these evaluations cover a lot, there are still many niche or one-off issues affecting thousands of files. A comprehensive list of everything would not be reasonable to create or maintain, but a list of some of these are provided in the supplemental materials.

Based on the evaluations presented here, we have a set of recommendations for MW and the metabolomics field. Actually, all of the recommendations in our last mwtab paper [6] still hold true now. The first major recommendation is to utilize versioning in the analysis files. Both the format requirements need to be versioned and the file contents need to be versioned. While the analysis files include a version key-value pair, it has not been effectively utilized and all analysis files still list a 1.0 version. Also, the mwTab specification documents for each version should be made available on the MW website. Currently only the latest version of the mwTab specifications description is available. Without format versioning, it is impossible to apply appropriate validations to analysis files that are following the format requirements at the time of their deposition. We brought the mwtab package up-to-date with the latest specification version available, but it was impossible to know when exactly some of the changes to the specification occurred since we could not look at the history changes to the specification. This made it harder to analyze which datasets were failing validation due to new requirements that are unfair to subject them to since they did not exist at the time of deposition.

The second major recommendation is to standardize/harmonize column names in the METABOLITES table. Without standardization and/or harmonization, reuse of large numbers of analysis files is pragmatically impossible. As a stop-gap, we have developed some tools that are now available in the new version of the mwtab package to help with the standardization/harmonization of column names in the METABOLITES table. The page in our documentation detailing how to use these tools is at https://moseleybioinformaticslab.github.io/mwtab/metadata_column_matching.html. These tools aren’t specific to metabolite data and could be used for any tabular data, but the prebuilt column finders are all specific to metabolite data and built from the datasets in MW.

The third major recommendation is to adopt UTF-8 character encoding in all mwTab and JSON formatted analysis files and in everything returned through MW’s REST interface. Such an adoption will prevent a variety of parsing errors. We also mentioned in our previous paper that a certification system certifying metadata quality and consistency could incentivize better depositions from depositors, and we still believe this to be true, but would recommend focusing on cleaning and improving the data and metadata that is already in MW first. However, such improvements should include a versioning of the contents of the analysis files.

## Supplementary Materials

Table S1: List of analysis IDs and the format of files that have parsing errors when trying to read them using the new version of the mwtab package, Table S2: List of analysis IDs and the format of files that have parsing errors when trying to read them using the previous version of the mwtab package, Table S3: List of analysis IDs and the format of files that have validation bugs when trying to validate them using the previous version of the mwtab package, and a list of many small issues found in MW files not mentioned in the paper.

## Author Contributions

Conceptualization, P.T.T. and H.N.B.M.; methodology, P.T.T. and H.N.B.M.; software, P.T.T.; validation, P.T.T.; formal analysis, P.T.T.; investigation, P.T.T.; resources, P.T.T. and H.N.B.M.; data curation, P.T.T.; writing—original draft preparation, P.T.T.; writing—review and editing, P.T.T. and H.N.B.M.; visualization, P.T.T.; supervision, H.N.B.M.; project administration, P.T.T. and H.N.B.M.; funding acquisition, H.N.B.M. Both authors have read and agreed to the published version of the manuscript.

## Institutional Review Board Statement

Not applicable.

## Informed Consent Statement

Not applicable.

## Data Availability Statement

The downloaded files from MW along with Python scripts for generating the results used in this paper are available in a Figshare item at https://doi.org/10.6084/m9.figshare.30924581.

## Acknowledgments

The authors would like to acknowledge the diligence that Shankar Subramaniam, Eoin Fahy, and the whole MW/UC San Diego team have put into provisioning FAIR access to metabolite studies and their incredible effort in keeping up with the exponential growth of the repository.

## Abbreviations

The following abbreviations are used in this manuscript:

MW: Metabolomics Workbench
MS: Mass Spectroscopy
NMR: Nuclear Magnetic Resonance
FAIR: Findable Accessible Interoperable Reusable
BSD: Berkeley Software Distribution
REST: Representational State Transfer
API: Application Programming Interface
FAIR: Findable Accessible Interoperable Reusable
CLI: Command Line Interface
JSON: JavaScript Object Notation
CSS: Cascading Style Sheets
PYPI: Python Package Index

## Supplemental Material

**Table S1.**
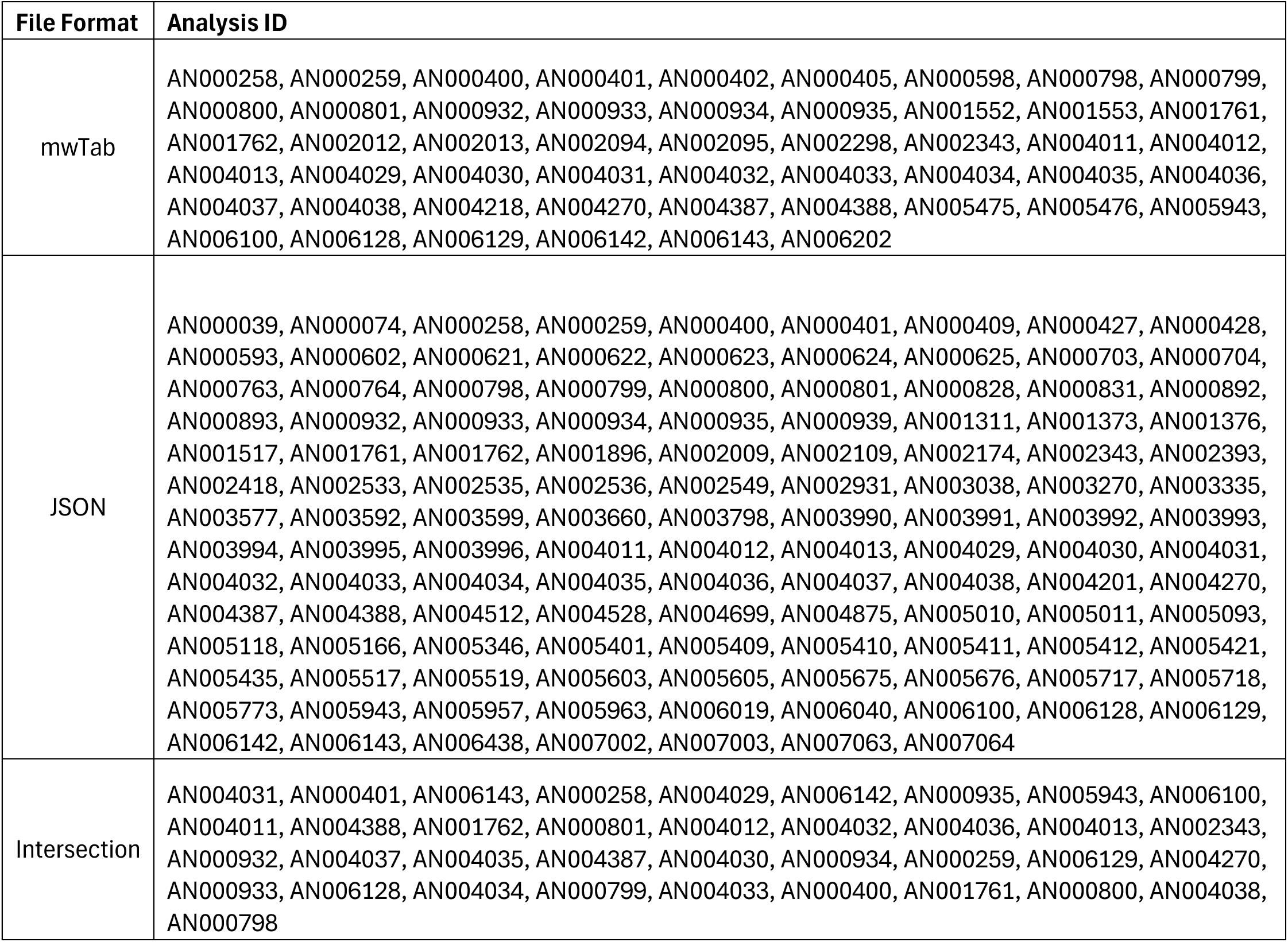
List of analysis IDs and the format of files that have parsing errors when trying to read them using the version 2.0.0 of the mwtab package.

| File Format | Analysis ID |
| --- | --- |
| mwTab | AN000258, AN000259, AN000400, AN000401, AN000402, AN000405, AN000598, AN000798, AN000799, AN000800, AN000801, AN000932, AN000933, AN000934, AN000935, AN001552, AN001553, AN001761, AN001762, AN002012, AN002013, AN002094, AN002095, AN002298, AN002343, AN004011, AN004012, AN004013, AN004029, AN004030, AN004031, AN004032, AN004033, AN004034, AN004035, AN004036, AN004037, AN004038, AN004218, AN004270, AN004387, AN004388, AN005475, AN005476, AN005943, AN006100, AN006128, AN006129, AN006142, AN006143, AN006202 |
| JSON | AN000039, AN000074, AN000258, AN000259, AN000400, AN000401, AN000409, AN000427, AN000428, AN000593, AN000602, AN000621, AN000622, AN000623, AN000624, AN000625, AN000703, AN000704, AN000763, AN000764, AN000798, AN000799, AN000800, AN000801, AN000828, AN000831, AN000892, AN000893, AN000932, AN000933, AN000934, AN000935, AN000939, AN001311, AN001373, AN001376, AN001517, AN001761, AN001762, AN001896, AN002009, AN002109, AN002174, AN002343, AN002393, AN002418, AN002533, AN002535, AN002536, AN002549, AN002931, AN003038, AN003270, AN003335, AN003577, AN003592, AN003599, AN003660, AN003798, AN003990, AN003991, AN003992, AN003993, AN003994, AN003995, AN003996, AN004011, AN004012, AN004013, AN004029, AN004030, AN004031, AN004032, AN004033, AN004034, AN004035, AN004036, AN004037, AN004038, AN004201, AN004270, AN004387, AN004388, AN004512, AN004528, AN004699, AN004875, AN005010, AN005011, AN005093, AN005118, AN005166, AN005346, AN005401, AN005409, AN005410, AN005411, AN005412, AN005421, AN005435, AN005517, AN005519, AN005603, AN005605, AN005675, AN005676, AN005717, AN005718, AN005773, AN005943, AN005957, AN005963, AN006019, AN006040, AN006100, AN006128, AN006129, AN006142, AN006143, AN006438, AN007002, AN007003, AN007063, AN007064 |
| Intersection | AN004031, AN000401, AN006143, AN000258, AN004029, AN006142, AN000935, AN005943, AN006100, AN004011, AN004388, AN001762, AN000801, AN004012, AN004032, AN004036, AN004013, AN002343, AN000932, AN004037, AN004035, AN004387, AN004030, AN000934, AN000259, AN006129, AN004270, AN000933, AN006128, AN004034, AN000799, AN004033, AN000400, AN001761, AN000800, AN004038, AN000798 |

**Table S2.**
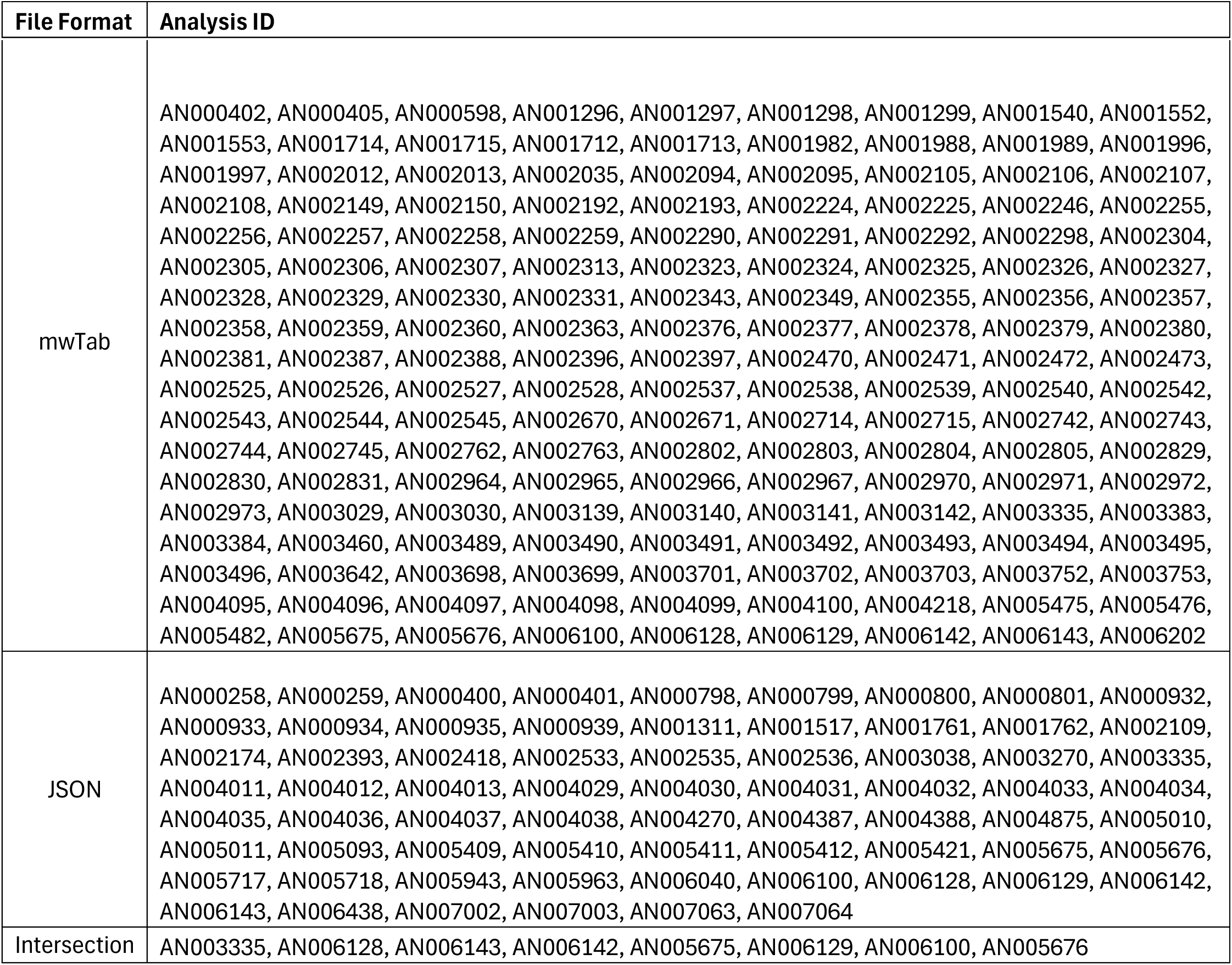
List of analysis IDs and the format of files that have parsing errors when trying to read them using version 1.2.5 of the mwtab package.

| File Format | Analysis ID |
| --- | --- |
| mwTab | AN000402, AN000405, AN000598, AN001296, AN001297, AN001298, AN001299, AN001540, AN001552, AN001553, AN001714, AN001715, AN001712, AN001713, AN001982, AN001988, AN001989, AN001996, AN001997, AN002012, AN002013, AN002035, AN002094, AN002095, AN002105, AN002106, AN002107, AN002108, AN002149, AN002150, AN002192, AN002193, AN002224, AN002225, AN002246, AN002255, AN002256, AN002257, AN002258, AN002259, AN002290, AN002291, AN002292, AN002298, AN002304, AN002305, AN002306, AN002307, AN002313, AN002323, AN002324, AN002325, AN002326, AN002327, AN002328, AN002329, AN002330, AN002331, AN002343, AN002349, AN002355, AN002356, AN002357, AN002358, AN002359, AN002360, AN002363, AN002376, AN002377, AN002378, AN002379, AN002380, AN002381, AN002387, AN002388, AN002396, AN002397, AN002470, AN002471, AN002472, AN002473, AN002525, AN002526, AN002527, AN002528, AN002537, AN002538, AN002539, AN002540, AN002542, AN002543, AN002544, AN002545, AN002670, AN002671, AN002714, AN002715, AN002742, AN002743, AN002744, AN002745, AN002762, AN002763, AN002802, AN002803, AN002804, AN002805, AN002829, AN002830, AN002831, AN002964, AN002965, AN002966, AN002967, AN002970, AN002971, AN002972, AN002973, AN003029, AN003030, AN003139, AN003140, AN003141, AN003142, AN003335, AN003383, AN003384, AN003460, AN003489, AN003490, AN003491, AN003492, AN003493, AN003494, AN003495, AN003496, AN003642, AN003698, AN003699, AN003701, AN003702, AN003703, AN003752, AN003753, AN004095, AN004096, AN004097, AN004098, AN004099, AN004100, AN004218, AN005475, AN005476, AN005482, AN005675, AN005676, AN006100, AN006128, AN006129, AN006142, AN006143, AN006202 |
| JSON | AN000258, AN000259, AN000400, AN000401, AN000798, AN000799, AN000800, AN000801, AN000932, AN000933, AN000934, AN000935, AN000939, AN001311, AN001517, AN001761, AN001762, AN002109, AN002174, AN002393, AN002418, AN002533, AN002535, AN002536, AN003038, AN003270, AN003335, AN004011, AN004012, AN004013, AN004029, AN004030, AN004031, AN004032, AN004033, AN004034, AN004035, AN004036, AN004037, AN004038, AN004270, AN004387, AN004388, AN004875, AN005010, AN005011, AN005093, AN005409, AN005410, AN005411, AN005412, AN005421, AN005675, AN005676, AN005717, AN005718, AN005943, AN005963, AN006040, AN006100, AN006128, AN006129, AN006142, AN006143, AN006438, AN007002, AN007003, AN007063, AN007064 |
| Intersection | AN003335, AN006128, AN006143, AN006142, AN005675, AN006129, AN006100, AN005676 |

**Table S3.**
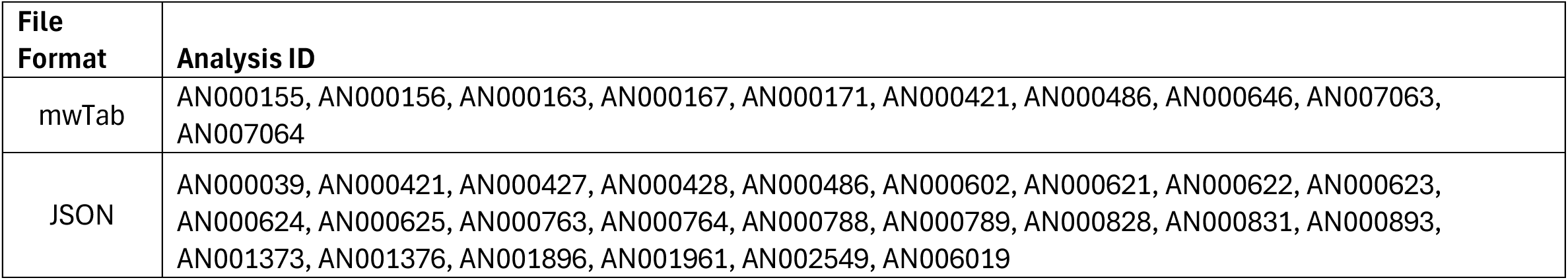
List of analysis IDs and the format of files that have validation bugs when trying to validate them using version 1.2.5 of the mwtab package.

| File Format | Analysis ID |
| --- | --- |
| mwTab | AN000155, AN000156, AN000163, AN000167, AN000171, AN000421, AN000486, AN000646, AN007063, AN007064 |
| JSON | AN000039, AN000421, AN000427, AN000428, AN000486, AN000602, AN000621, AN000622, AN000623, AN000624, AN000625, AN000763, AN000764, AN000788, AN000789, AN000828, AN000831, AN000893, AN001373, AN001376, AN001896, AN001961, AN002549, AN006019 |

The following is simply an unorganized list of many issues to do with mwTab files usually along with the analysis ID of a file exhibiting that issue:

- Too many elements in the header. There are random filenames/strings mixed in with the key value pairs as well as duplicate keys. AN000037
- Additional “Samples”, “Factors”, and “metabolite_name” lines that aren’t where they are supposed to be or in completely incorrect sections.
- Incorrectly labeled “Samples” sections. For example, a “Samples” line instead having “metabolite_name”.
- Slight variations on list headers, such as “Metabolite Name” instead of “metabolite_name”. AN000144
- Spacing out of specification. CREATED_ON is biggest offender, but other blocks also have the incorrect spaces according to the specification. AN000086
- Extra tabs at the end of some lines. Mostly on list header lines, “Factors”, “Samples”, “metabolite_name”. AN000152
- Extra spaces in SUBJECT_SAMPLE_FACTORS. AN000152
- Incorrect prefix. For example, MS:INSTRUMENT_NAME instead of CH:INSTRUMENT_NAME in the #CHROMATOGRAPHY section. AN000037
- Unknown section name. For example, FACTORS. AN004560
- RESULTS_FILE in the wrong block. AN002939
- Extra empty key value pairs in JSON tables. There is an extra ““:“” at the end of every dictionary. AN000020
- Values in the #METABOLITES block have more values than headers (this could happen in other blocks). Seems to stem from values that have a tab in the value, for example “\t65359”. AN005166 AN003577 AN003592 AN003599 AN003660 AN003798 AN004512 AN005118
- Values in the #METABOLITES block have more values than headers (this could happen in other blocks. Similar to the one above, except in this case there are just too many values and not just extra tabs. AN002931 AN002681 AN002682 AN002683 AN002682 AN004201 AN004528 AN004987
- ’#’ at the beginning of lines that aren’t new blocks, for example #MS:MS_RESULTS_FILE. AN004368
- Sub-sections are repeated, such as, “MS:INSTRUMENT_NAME Agilent 6220 TOF” appearing twice, but not on consecutive lines. Sometimes on consecutive lines. Most are 1 line repeated directly under, and most are in MS section. AN001532 AN001449
- Duplicate factor names in SUBJECT_SAMPLE_FACTORS. AN000379
- Duplicate keys in Additional sample data, but the order matters on the keys. AN001558
- Extra “#END” line at the end of the file. AN001859
- The two-letter code at the beginning of the line is not correct. Example: “_1 CH:CHROMATOGRAPHY” AN002702
- missing the two-letter code, “NM:”, at the start of the RESULTS_FILE line. AN002939 AN004560
- Additional data keys in SUBJECT_SAMPLE_FACTORS with too many ‘=’. AN000258 AN000259
- Factor keys in SUBJECT_SAMPLE_FACTORS with too many ‘:’. AN000400 AN000401
- Newline that shouldn’t be there. For example, an email that has a newline just before the @ part. AN000402 AN000598 AN002012 AN002013 AN002094 AN002095 AN004218
- Missing a tab after spaces and before payload. For example, “PR:PHONE --” has no tab before “--”. AN000405
- Every NMR_BINNED_DATA set has malformed JSON without a ‘NMR_BINNED_DATA’ key and instead just has a ‘Data’ key. It should be under a ‘NMR_BINNED_DATA’ key just like all the other data sets. This is a recent change. Older downloaded versions had NMR_BINNED_DATA sections just as expected. AN000041
- Many data sets, around 250, have an extra blank line between all of the lines. AN000001
- EXTENDED sections don’t appear in JSON version of the file.
- The full data is copied twice inside the file. Some variations of this have all of the section names copied in twice but with no sub-sections, sort of like a skeleton of a file. The skeletons are sometimes before the actual data and sometimes after. AN005718
- Badly formed results file line and line outside of MS/NMR section. AN001881

